# A Dual Assay to Compare Protein Levels and Toxicity of Alpha-Synuclein Variants: Acute Expression of Wild-Type versus S129A

**DOI:** 10.1101/2025.09.23.678012

**Authors:** Baoyi Li, Oren A. Levy, Nagendran Ramalingam, Ulf Dettmer

## Abstract

Parkinson’s disease (PD) affects over 12 million people worldwide and has the fastest-growing global impact. The pathological hallmark is the presence of Lewy bodies and Lewy neurites, which are intraneuronal lesions enriched in aggregated alpha-synuclein (αS) that is typically hyper-phosphorylated at serine 129. Therefore, lowering phosphoserine 129 (pS129) may be a viable therapeutic strategy to treat PD. However, pS129 has also been proposed to regulate synaptic transmission and αS degradation. In both cases, inhibiting pS129 could be detrimental. Here, we developed a sensitive assay in a human neuroblastoma model and utilized it to assess the relative expression levels and cytotoxicity of pS129 by comparing αS wild-type (WT) vs. S129A (pS129-deficient). We show that the S129A mutant does not affect the acute expression levels or toxicity of αS in a transient transfection paradigm. This provides new insight into the intricate interplay between αS, phosphorylation, toxicity, and degradation. Our assay provides a versatile platform for understanding disease-relevant mechanisms and opens novel avenues for the design of future therapeutic interventions in PD and other α-synucleinopathies.

## Introduction

Parkinson’s disease (PD) is the second most common neurodegenerative condition, affecting over 12 million people worldwide and has the fastest-growing global impact of all neurodegenerative conditions. Over the past generation, its global burden has more than doubled, driven mainly by an aging population (GBD 2016 Neurology Collaborators, 2019). It is estimated that in 2050 there will be 25.2 million people with PD around the world, up from 11.9 million people in 2021 (Su et al., 2025). PD affects the quality of life of people who suffer from it, but current treatments are only able to manage PD symptoms rather than stop disease progression. Thus, research into therapies to slow or stop PD remains essential.

The neuronal protein alpha-synuclein (αS) was linked to PD pathology when it was identified in intraneuronal aggregates in PD brains (Spillantini et al., 1997). αS is a small, 140-amino acid polypeptide encoded by the *SNCA* gene and is mainly present at the presynaptic terminals of neurons, where it has been implicated in regulating dopamine reuptake and release, membrane curvature, and vesicle trafficking (reviewed by Bendor et al., 2013). It exists *in vivo* in a tightly regulated equilibrium between the cytosol and transiently bound to membranes (reviewed by Yeboah et al., 2019). Excess interactions between αS and cell membranes may play a key role in αS aggregation (reviewed by Li & Dettmer, 2024). *SNCA* multiplications (Chartier-Harlin et al., 2004) and missense mutations such as A53T (Polymeropoulos et al., 1997) and E46K (Zarranz et al., 2004) have been implicated in familial PD. Therefore, αS is also genetically linked with the condition. In the hallmark pathological lesions of PD and other α-synucleinopathies, αS is abundantly phosphorylated, and around 90% of the aggregated αS may be modified by phosphorylation at serine 129 (pS129), while only a small fraction (<4%) of αS is phosphorylated under physiological conditions (Anderson et al., 2006; Fujiwara et al., 2002; Neumann et al., 2002). This has led to the widely cited conclusion that pS129 is linked to PD pathology. However, the reality seems to be more nuanced.

Many previous studies have ignored the normal role of pS129. Indeed, phosphorylation at S129 may be a normal event in mouse brain or adult human brain (Hirai et al., 2004; Muntané et al., 2012). Our recent work showed that neuronal activity increases pS129 in cultured neurons, which in turn positively regulates synaptic transmission (Ramalingam et al., 2023). It has also been suggested that pS129 may even have neuroprotective effects by augmenting the degradation of aggregated αS in vivo (Oueslati et al., 2013). Thus, the detailed roles and effects of this post-translational modification in αS physiology and PD pathogenesis will require further research.

To deepen our understanding of αS physiology and pathophysiology, including the role of pS129, sensitive and robust assays for quantifying αS levels as well as αS-related toxicity are critical. In this context, internal ribosome entry site (IRES)-based bicistronic constructs have been used in other systems since their first discovery in viral RNAs (Jang et al., 1988; Pelletier & Sonenberg, 1988) for the co-expression of multiple proteins from a single transcript. For example, IRES elements have enabled co-expression of a non-tagged chaperone and a fluorescent reporter (San Gil et al., 2017). To our knowledge, however, no prior study has applied this strategy to compare the effects of missense mutations on αS expression levels and αS-associated toxicity in parallel. Our αS-IRES-mCherry strategy thus represents a novel adaptation tailored for dynamic, quantitative monitoring of αS expression and cytotoxicity.

Using this approach, we investigated the role of S129 phosphorylation by comparing WT αS with the phospho-deficient S129A mutant. Our results show that S129A expression did not alter αS levels or increase toxicity in neuroblastoma cells under the conditions tested, suggesting that the absence of phosphorylation at this site does not confer acute toxicity.

## Materials and Methods

All materials mentioned were purchased from Invitrogen unless stated otherwise.

### Cell culture

BE (2)-M17 human neuroblastoma (M17D) cells were chosen as they have been shown to have rapid growth *in vitro*, and improved reproducibility of experiments compared to other models such as SH-SY5Y cells (Carvajal-Oliveros et al. 2024). M17D cells were cultured in Dulbecco’s modified Eagle’s medium (DMEM) supplemented with 10% fetal bovine serum (Sigma) and 2 mM L-glutamine in a 5% CO_2_ atmosphere at 37 ºC. Cells plated in 48-well plates were transfected using Lipofectamine 2000 according to the manufacturer’s directions. Cells were harvested 24 h, 48 h, and 72 h post transfection.

### cDNA cloning

αS mutants were cloned into the pLVX-IRES-mCherry plasmid (Clontech) by restriction-enzyme-based cloning using SpeI/NotI sites. To assess the functional role of phosphorylation at S129, a phospho-deficient S129A mutant was engineered by substituting the amino acid serine with alanine, thus preventing phosphorylation. We used the plasmids pLVX-IRES-mCherry/αS-WT (Dettmer et al., 2017) and pLVX-IRES-mCherry/αS-WT-S129A. See Supplementary Figure 1 for the amino-acid sequences of the αS variants.

### Cell lysis

M17D cells were collected from the plate 24 h, 48 h, and 72 h post-transfection. Cells were centrifuged at 3,000 *g* for 2 min at room temperature. Culture media was aspirated, and the pellet was resuspended in PBS. Next, cells were centrifuged at 3,000 *g* (room temperature; 2 min) and PBS was aspirated. Cells were lysed on ice using 75 μL of 0.5% Triton X-100 with phosphatase and protease inhibitors. This was followed by centrifugation at 16,100 *g* for 15 min at 4 ºC. 15 μL of 4x protein sample loading buffer (Li-Cor) was added to new 1.5 mL tubes and 45 μL of the supernatant was added. Samples were boiled for 3 min and centrifuged at 3,000 *g* for 2 min.

### Immunoblotting

12 μL of lysate was loaded per lane. Samples were electrophoresed on NuPAGE 4– 12% Bis-Tris gels with NuPAGE MES-SDS running buffer and SeeBlue Plus2 marker. Gels were electroblotted onto nitrocellulose membranes using an iBlot 2NC stack and iBlot 2 device (20 V for 1 min, 23 V for 4 min, then 25 V for 2 min). Membranes were incubated in 0.4% paraformaldehyde for 20 min, rinsed with TBS, stained with 0.1% Ponceau S in 5% acetic acid, rinsed with water and blocked in the blocking buffer (Li-Cor blocking buffer ‘TBS’) for 30 min at room temperature. Membranes were incubated in primary antibodies in the blocking buffer either for 1 h at room temperature or overnight at 4 ºC. Membranes were washed 3 × 10 min in TBST or PBST at room temperature and incubated in secondary antibodies in the blocking buffer for 40 min at room temperature. Membranes were washed 3 × 10 min in TBST or PBST and scanned (Odyssey CLx, Li-Cor).

### Antibodies

Primary antibodies used in Western blotting were monoclonals 15G7 to αS (Enzo Life Sciences; 1:250 in Western blot), D1R1R to pS129 (Cell Signaling; 1:2,000 in Western blot), AB9484 to GAPDH (Abcam; 1:3,000 in Western blot), and 16D7 to mCherry (Thermo Scientific; 1:300 in Western blot). Secondary antibodies used were polyclonal Li-Cor IRDye anti-mouse, anti-rabbit, and anti-rat (926-68070, 926-68071, and 926-32219 respectively; 1:10,000 in Western blot).

### Statistical analyses

We performed unpaired two-tailed t-tests to compare WT and S129A αS using GraphPad Prism version 10 following the program’s guidelines. Normal (Gaussian) distribution was confirmed for all values in bar graphs by Shapiro–Wilk and Kolmogorov–Smirnov tests. Graphs represent means ± standard deviation (SD).

### Live-cell Imaging

Live-cell imaging was performed using the IncuCyte S3-C2 system (Sartorius). Images from phase-contrast and red fluorescence channels were acquired every 4 h using a 10× objective. Red fluorescence integrated intensity and phase-channel cell confluence in % were quantified using the program’s standard internal “processing definitions”. Data from different wells of a 96-well plate (biological replicates) were averaged, and 3 independent experiments were performed on 3 different days, followed by analysis via GraphPad Prism as described above. Results are presented as mean ± standard deviation (SD).

## Results

### Establishing a robust assay in neuroblastoma cells

We aimed to develop a robust cellular assay with two distinct readouts: (1) detection and relative quantification of protein levels of different αS variants, and (2) assessment of their relative cytotoxicity.

We utilized cDNA plasmids encoding αS and mCherry as a bicistronic construct, separated by an internal ribosomal entry site (IRES). The two genes are under the same promoter and transcribed into the same mRNA, thus they are co-expressed by cells. This way, Western blot for mCherry serves as an intrinsic control for transfection efficiency. In addition, monitoring mCherry fluorescence signals in live cells serves as a proxy for αS-related toxicity using live-cell fluorescence microscopy through Incucyte. If the production of an αS variant is toxic, this would lead to a decrease in proliferation and increase in cell death, which would result in a reduction in red mCherry fluorescence, enabling real-time toxicity assessment using Incucyte (Figure 1).

**Figure 1:**
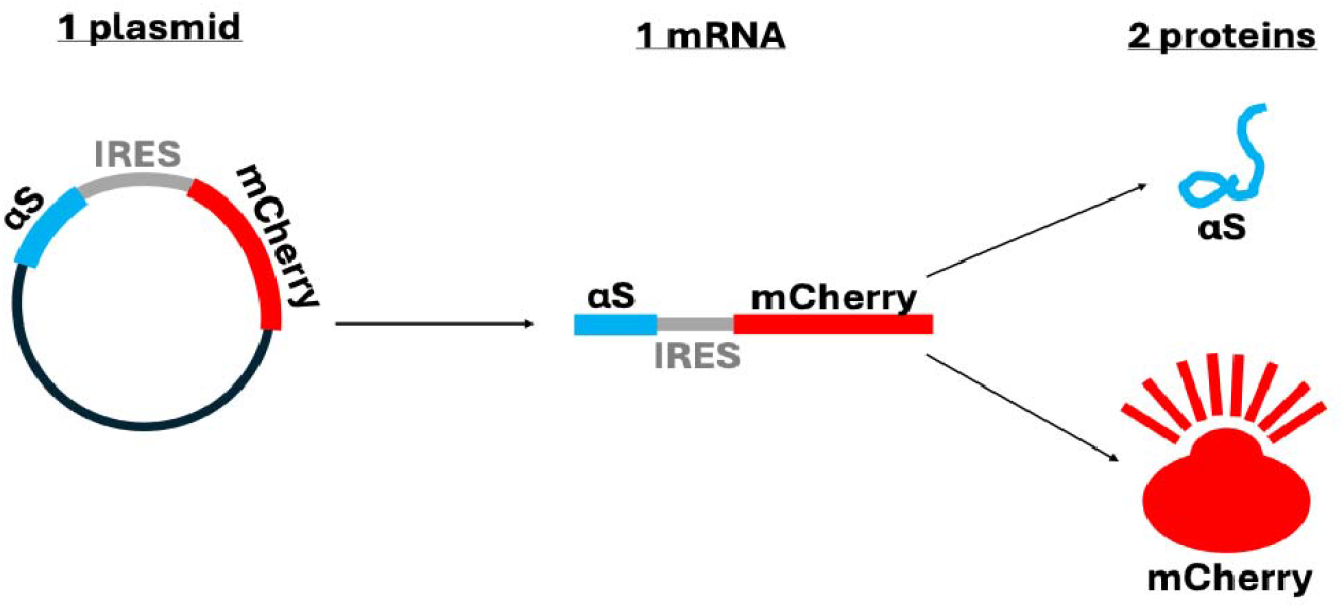
Schematic for cDNA plasmid. Plasmid encodes αS and mCherry, separated by IRES, as a bicistronic construct. One plasmid leads to the formation of one mRNA to produce two proteins, αS and the red fluorescent reporter protein mCherry.

We first identified a set of conditions that produced reliable and interpretable results. For Western blot analysis of αS levels, we plated cells in 48-well plates in such a way that they would reach 50-80% confluence on the day of transfection. For transfection, we used Lipofectamine 2000 according to the manufacturer’s protocol. As a highly suitable primary antibody combination, we identified monoclonal rat antibody 15G7 to αS, monoclonal rabbit antibody D1R1R to pS129, monoclonal mouse antibody EPR6256 to GAPDH, and monoclonal rabbit antibody 16D7 to mCherry. Secondary antibodies were polyclonal Li-Cor IRDye anti-mouse, anti-rabbit, and anti-rat (926-68070, 926-68071, and 926-32219 respectively). These conditions made for a robust, sensitive assay enabling the assessment of protein levels and toxicity without artifacts based on transfection efficiency. To assess cell viability, we transfected ∼50% confluent M71D cells in 96-well plates, followed by Incucyte analysis of mCherry and bright field signals for 5 days.

### Incucyte live-cell microscopy confirms transfection via mCherry expression

To assess the transfection of our M17D neuroblastoma cell cultures, we visualized the red mCherry signal by live-cell fluorescence microscopy using the Incucyte imaging system 24 h post-transfection. As expected, no red signals were detected in control-transfected M17D neuroblastoma cells, while mCherry red was detected for those cells transfected with αS WT and S129A constructs (Figure 2). An estimated >90% of the cells displayed visible levels of mCherry signal.

**Figure 2:**
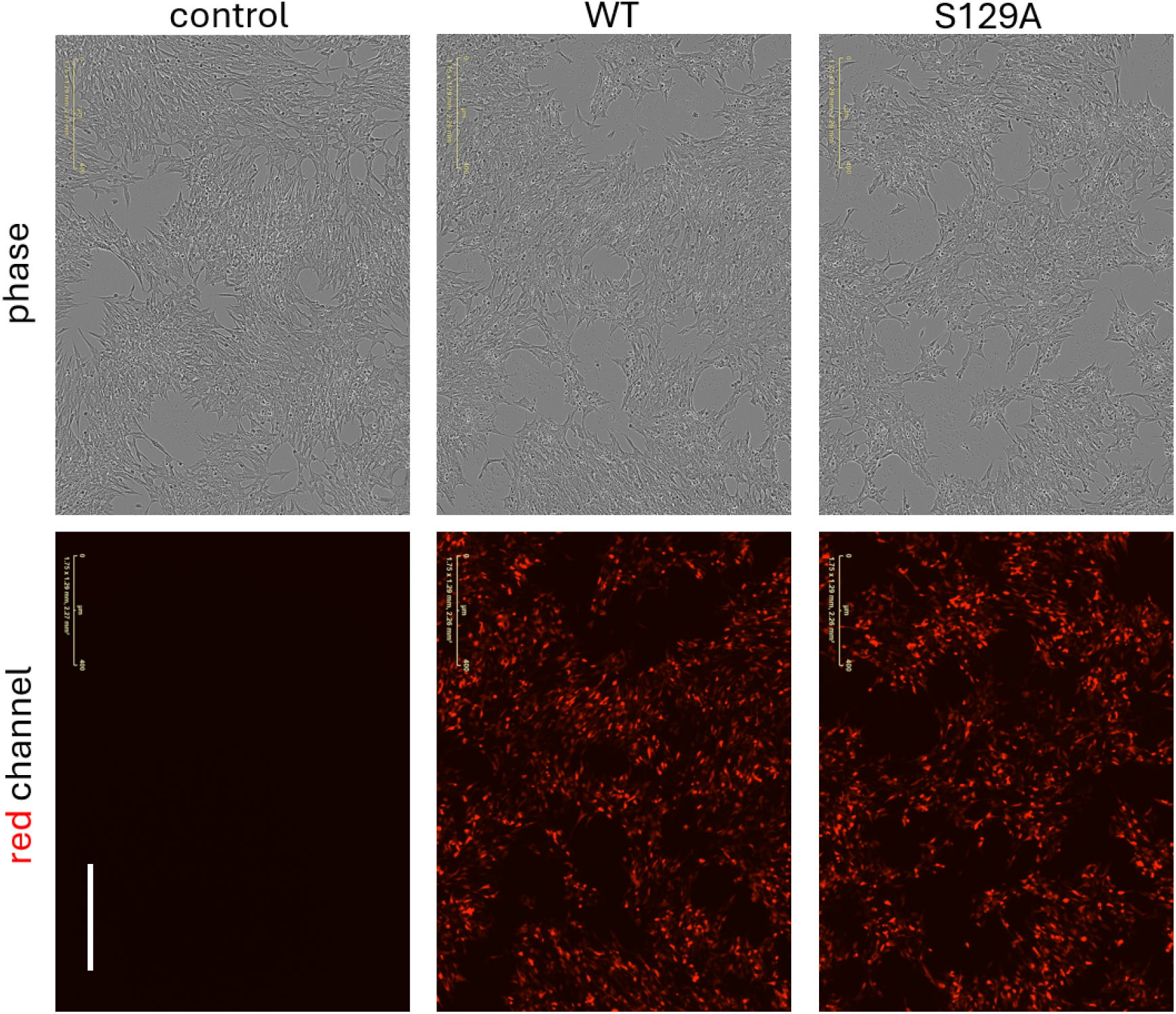
Successful transfection of cells confirmed by Incucyte. Equally scaled, representative epifluorescence images of M17D neuroblastoma cells transfected with plasmids expressing αS (WT or S129A) and the red fluorescent mCherry reporter (control: mock transfection without plasmid). Phase channel displays M17D cells without fluorescence. Red channel shows mCherry signal as a proxy for transfection and cell health. Scale bar represents 400 μm.

### Levels of WT vs. S129A αS

To investigate the effect of S129A on WT αS, M17D cells were transfected mock (-), αS WT, and αS S129A. We then measured the levels of mCherry, GAPDH (loading control), total αS, and pS129 αS by Western blotting (Figure 3). The pS129-specific antibody detected WT but not S129A, confirming the identity of the latter. Moreover, there were no significant differences in WT vs. S129A total αS levels (normalized to mCherry) for three days after transfection. We also observed that the mCherry signal forms a doublet for all αS mutants (Fig. 3). This may be explained by the presence of an alternative translation initiation site in the mCherry coding sequence (Fages-Lartaud et al., 2022). Ribosomes can start translation here, producing a truncated protein isoform (around 1-2 kDa smaller). Both isoforms are stable, fluorescent, and antibody-reactive, and both signals were combined in our analyses.

**Figure 3:**
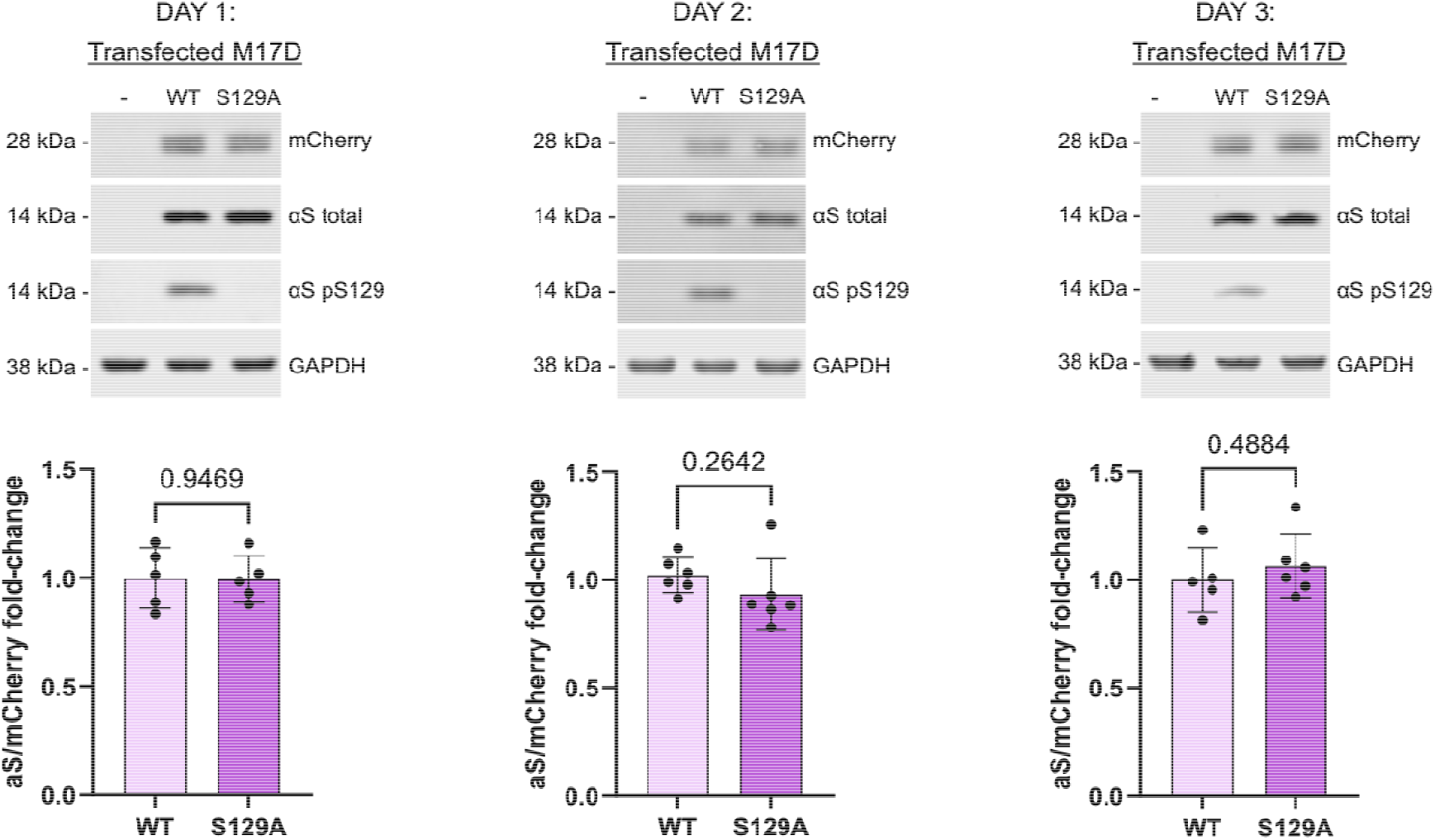
Levels of WT and S129A αS. Western blot for mCherry, total αS, pS129 αS, and GAPDH. Quantification of total αS to WT (normalized to mCherry) for M17D cells transfected with WT or S129A variants. Cells were harvested 24 h, 48 h, and 72 h post-transfection. Graphs represent *N* = 3 independent experiments done in duplicates. Each data point represents 1 biological replicate. Mean ± SD. Normal distribution was confirmed for all values in bar graphs by Shapiro–Wilk and Kolmogorov–Smirnov tests. Unpaired two-tailed t-tests; no differences were statistically significant (threshold *p* < 0.05).

### Toxicity of WT vs. S129A αS

Furthermore, we assessed whether S129A affects the toxicity of αS. M17D cells were transfected αS WT and αS S129A, and we used live-cell fluorescence microscopy with the Incucyte system to measure the mCherry fluorescence signal over a period of 5 days. Because αS and mCherry were co-expressed bicistronically in the cDNA plasmids, mCherry fluorescence served as a real-time proxy of αS-related toxicity. In 3 independent experiments, S129A did not have a significant effect on mCherry fluorescence signal, and thus the toxicity of αS (Figure 4).

**Figure 4:**
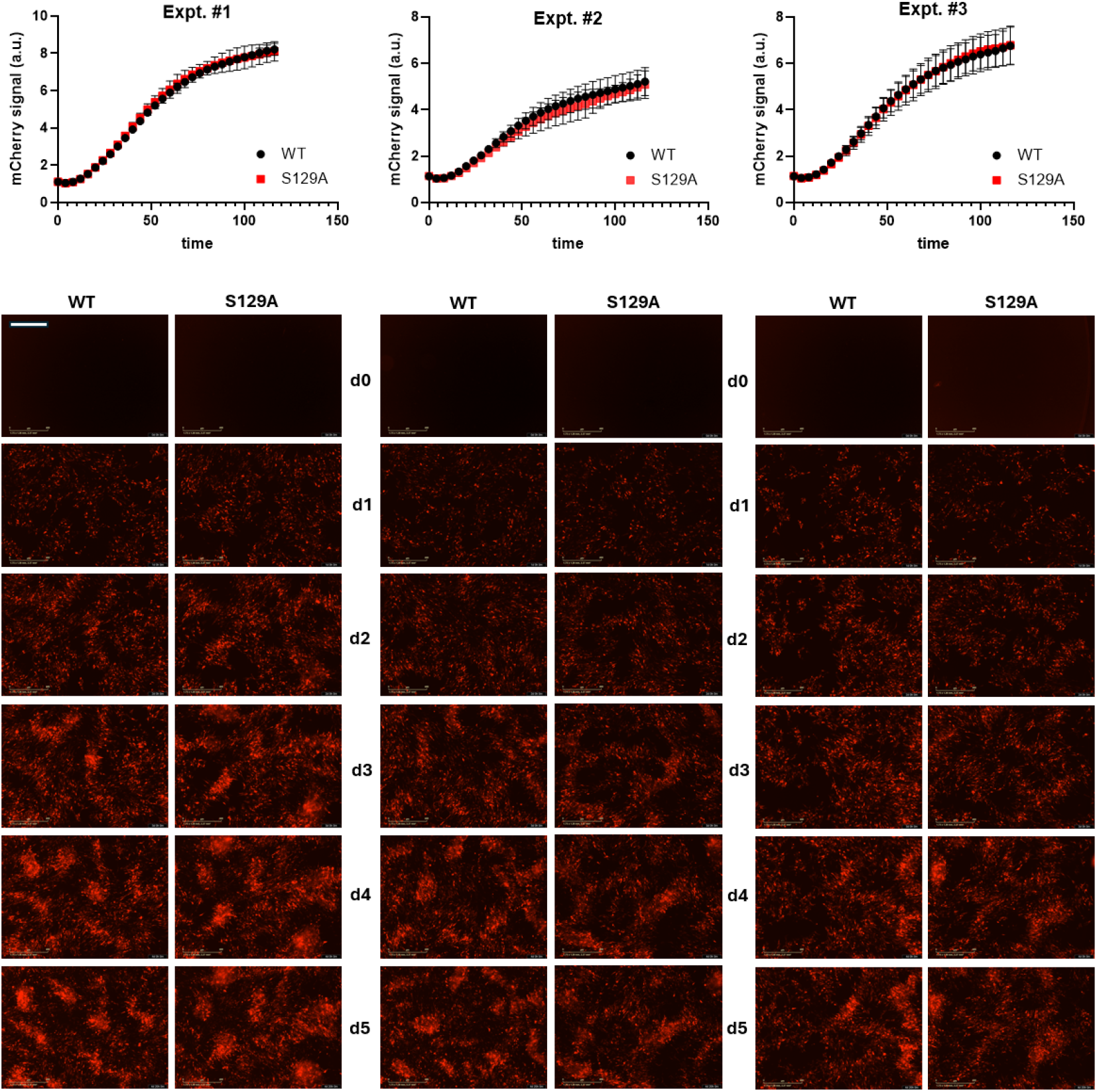
Toxicity of WT and S129A αS. Time-course quantification of mCherry fluorescence signal (arbitrary units, a.u.), normalized to control, measured with Incucyte in M17D cells expressing WT or S129A αS. Time points indicate hours post-transfection. Panels A-C represent *N* = 3 independent experiments over 5 days. Each time point represents *n* = 5 biological replicates. d0 = day 0 (pre-transfection), d1 = day 1 after transfection, etc. Mean ± SD. Scale bar, 200 µm.

## Discussion

This study presents a dual-readout assay designed to quantify αS expression and assess associated toxicity in a human neuroblastoma model. The approach was applied to compare human αS WT to pS129-deficient S129A. The approach integrates a bicistronic αS-IRES-mCherry construct with live-cell imaging, enabling simultaneous measurement of protein levels and cell viability over time. The use of mCherry as an internal control for transfection efficiency and cell health provides a robust normalization strategy, reducing variability and enhancing interpretability. Bicistronic expression systems using IRES elements are well-established in research (Jang et al., 1988; Pelletier & Sonenberg, 1988). For example, they have been employed to co-express untagged proteins with fluorescent reporters (San Gil et al., 2017). However, to our knowledge, this is the first implementation specifically pairing αS with mCherry via an IRES for time-resolved quantification of both expression and toxicity. Compared to conventional approaches such as endpoint viability assays or Western blotting without an intrinsic control from bicistronic protein translation, our strategy offers several advantages. The use of live-cell imaging provides temporal resolution that is lacking in static assays, while the internal mCherry control enables normalization across conditions and replicates. These features contribute to improved sensitivity and reproducibility. Certain limitations of IRES-based bicistronic systems must be considered, even though they should not have affected our outcomes. Expression from the downstream cistron is typically weaker and more variable, and the efficiency of IRES-driven translation can depend on cell type and physiological state (Hennecke et al., 2001; Mizuguchi et al., 2000). Alternatives such as 2A peptide systems (Kim et al., 2011; Szymczak et al., 2004) offer more stoichiometric co-expression, but also have drawbacks such as the presence of additional amino-acids in the protein of interest (“scar”).It should also be noted that mCherry half-life can be very long (Bell et al., 2025), which may mask certain effects when monitoring toxicity.

In our assay, the phospho-deficient S129A mutant did not alter αS levels or increase toxicity under the tested conditions (neuroblastoma cells, transient cDNA transfection). This absence of an effect provides a useful reference point for interpreting prior literature on the functional consequences of pS129. The broader literature on pS129 has revealed conflicting findings regarding its potential role in αS toxicity. Some studies suggest that pS129 promotes toxicity, e.g.: proteasome inhibition increased pS129 and was associated with neuronal cell death (Chau et al., 2009); phosphorylation was linked to distinct αS strain formation (Ma et al., 2016); and pS129 αS was reported to disrupt stress signaling by inhibiting polo⍰like kinase 2 (Wang et al., 2012). However, other studies argue against a direct toxic role: phosphorylation did not prevent the generation of toxic αS species in a rat model of PD (Azeredo da Silveira et al., 2009); S129 mutants did not alter nigrostriatal toxicity in a rat model of PD (McFarland et al., 2009); and enhancing pS129 via overexpression of polo-like kinases 2 and 3 in nigral dopamine neurons was not detrimental to their survival and function (Buck et al., 2015). Together, these findings highlight an ongoing debate as to whether pS129 is pathogenic, protective, or neutral (Oueslati, 2016; Ramalingam et al., 2024; Ramalingam & Dettmer, 2023). Beyond toxicity, pS129 has also been implicated in αS turnover and degradation. In a yeast αS model, pS129 accelerated αS turnoverc (Tenreiro et al., 2014). In mammalian (overexpression) systems, polo-like kinase 2 overexpression enhanced autophagic clearance of αS and suppressed toxicity (Oueslati et al., 2013). Proteasome inhibition was shown to elevate pS129 without changing total αS levels (Chau et al., 2009; Machiya et al., 2010; Ramalingam, Brontesi, et al., 2023), and phosphorylated αS was targeted to the proteasome in a ubiquitin-independent manner (Machiya et al., 2010). Most recently, RXR agonist–mediated signaling was shown to stimulate lysosomal clearance of αS in patient-derived neurons, with evidence that polo⍰like kinase 2 and pS129 contributed to this beneficial effect (Tripathi et al., 2024). These studies collectively suggest that pS129 may influence αS degradation under certain circumstances (excess, aggregation, ectopic expression under certain promoters, among others).

Overall, the observation that S129A did not alter αS levels or toxicity in our neuroblastoma model should be interpreted with caution. This system provides a useful screening platform, but it only partially recapitulates mature neurons. In addition, transient cDNA transfection creates a non-steady-state condition: it leads to the immediate production of high levels of encoded proteins, but then the DNA gets diluted out and degraded in the dividing cultures, so transgene transcription drops again. Effects of S129A may differ in viral overexpression systems, primary cultures, iPSC-derived neurons, or *in vivo* models. Given the complex interplay of S129 phosphorylation with αS degradation, aggregation, and synaptic function (Ramalingam, Brontesi, et al., 2023; Ramalingam et al., 2024; Ramalingam, Jin, et al., 2023; Ramalingam & Dettmer, 2023), further work will be needed. Moving forward, this assay is flexible and adaptable. It can be adapted to study familial PD mutations (e.g., A53T, E46K), *SNCA* multiplications, and pharmacological modulators of αS expression and clearance. It could also be combined with iPSC-derived neuronal models or in vivo paradigms to provide deeper mechanistic insights.

In summary, the αS-IRES-mCherry assay described here provides a technically sound and adaptable platform for studying αS biology. It allows quantitative and dynamic assessment of protein expression and toxicity in living cells, certain caveats about the IRES approach (Hennecke et al., 2001; Mizuguchi et al., 2000) should not have affected our key finding that the S129A phospho-deficient mutant did not alter αS levels or toxicity under the chosen conditions of short-term expression in neuroblastoma cells.

## Acknowledgements

We thank A. Tripathi and H. Alnakhala for assisting with cell culture and data acquisition.

## Author contributions

B.L., O.L., N.R., U.D. designed the research; N.R., U.D. supervised the research; B.L., U.D. performed the experiments; O.L., N.R., U.D. generated tools; B.L., N.R., U.D. analyzed the data; B.L., U.D. visualized the data; B.L. wrote the original manuscript; B.L., O.L, N.R., U.D. reviewed and edited the manuscript.

## Notes

### Competing Interest Statement

The authors have declared no competing interest.

### Summary of Updates

The ORCID iD of author Baoyi Li was added.

## Bibliography

Anderson, J. P., Walker, D. E., Goldstein, J. M., De Laat, R., Banducci, K., Caccavello, R. J., Barbour, R., Huang, J., Kling, K., & Lee, M. (2006). Phosphorylation of Ser-129 is the dominant pathological modification of α-synuclein in familial and sporadic Lewy body disease. Journal of Biological Chemistry, 281 (40), 29739–29752.

Azeredo da Silveira, S., Schneider, B. L., Cifuentes-Diaz, C., Sage, D., Abbas-Terki, T., Iwatsubo, T., Unser, M., & Aebischer, P. (2009). Phosphorylation does not prompt, nor prevent, the formation of alpha-synuclein toxic species in a rat model of Parkinson’s disease. Human Molecular Genetics, 18(5), 872–887. 10.1093/hmg/ddn417

Bell, K. S., Ko, S., Ali, S., Bognar, B., Khmelkov, M., Rau, N., Peng, O. K., Eyuboglu, M., Paine, J., Tong, A., Saria, A., Agrawal, S., Davies, K. J. A., & Tower, J. (2025). Tissue-Specific Fluorescent Protein Turnover in Free-Moving Flies. Insects, 16(6), 583. 10.3390/insects16060583

Bendor, J. T., Logan, T. P., & Edwards, R. H. (2013). The function of α-synuclein. Neuron, 79(6), Article 6. 10.1016/j.neuron.2013.09.004

Buck, K., Landeck, N., Ulusoy, A., Majbour, N. K., El-Agnaf, O. M. A., & Kirik, D. (2015). Ser129 phosphorylation of endogenous α-synuclein induced by overexpression of polo-like kinases 2 and 3 in nigral dopamine neurons is not detrimental to their survival and function. Neurobiology of Disease, 78, 100–114. 10.1016/j.nbd.2015.03.008

Chartier-Harlin, M.-C., Kachergus, J., Roumier, C., Mouroux, V., Douay, X., Lincoln, S., Levecque, C., Larvor, L., Andrieux, J., & Hulihan, M. (2004). α-synuclein locus duplication as a cause of familial Parkinson ‘s disease. The Lancet, 364 (9440), 1167–1169.

Chau, K.-Y., Ching, H. L., Schapira, A. H. V., & Cooper, J. M. (2009). Relationship between alpha synuclein phosphorylation, proteasomal inhibition and cell death: Relevance to Parkinson’s disease pathogenesis. Journal of Neurochemistry, 110(3), Article 3. 10.1111/j.1471-4159.2009.06191.x

Dettmer, U., Ramalingam, N., von Saucken, V. E., Kim, T.-E., Newman, A. J., Terry-Kantor, E., Nuber, S., Ericsson, M., Fanning, S., Bartels, T., Lindquist, S., Levy, O. A., & Selkoe, D. (2017). Loss of native α-synuclein multimerization by strategically mutating its amphipathic helix causes abnormal vesicle interactions in neuronal cells. Human Molecular Genetics, 26(18), Article 18. 10.1093/hmg/ddx227

Fages-Lartaud, M., Tietze, L., Elie, F., Lale, R., & Hohmann-Marriott, M. F. (2022). mCherry contains a fluorescent protein isoform that interferes with its reporter function. Frontiers in Bioengineering and Biotechnology, 10, 892138.

Fujiwara, H., Hasegawa, M., Dohmae, N., Kawashima, A., Masliah, E., Goldberg, M. S., Shen, J., Takio, K., & Iwatsubo, T. (2002). Alpha-Synuclein is phosphorylated in synucleinopathy lesions. Nature Cell Biology, 4(2), Article 2. 10.1038/ncb748

GBD 2016 Neurology Collaborators. (2019). Global, regional, and national burden of neurological disorders, 1990-2016: A systematic analysis for the Global Burden of Disease Study 2016. The Lancet. Neurology, 18(5), 459–480. 10.1016/S1474-4422(18)30499-X

Hennecke, M., Kwissa, M., Metzger, K., Oumard, A., Kröger, A., Schirmbeck, R., Reimann, J., & Hauser, H. (2001). Composition and arrangement of genes define the strength of IRES-driven translation in bicistronic mRNAs. Nucleic Acids Research, 29(16), 3327–3334. 10.1093/nar/29.16.3327

Hirai, Y., Fujita, S. C., Iwatsubo, T., & Hasegawa, M. (2004). Phosphorylated α-synuclein in normal mouse brain. FEBS Letters, 572 (1–3), 227–232.

Jang, S. K., Kräusslich, H. G., Nicklin, M. J., Duke, G. M., Palmenberg, A. C., & Wimmer, E. (1988). A segment of the 5’ nontranslated region of encephalomyocarditis virus RNA directs internal entry of ribosomes during in vitro translation. Journal of Virology, 62(8), 2636–2643. 10.1128/JVI.62.8.2636-2643.1988

Kim, J. H., Lee, S.-R., Li, L.-H., Park, H.-J., Park, J.-H., Lee, K. Y., Kim, M.-K., Shin, B. A., & Choi, S.-Y. (2011). High cleavage efficiency of a 2A peptide derived from porcine teschovirus-1 in human cell lines, zebrafish and mice. PloS One, 6(4), e18556. 10.1371/journal.pone.0018556

Li, B., & Dettmer, U. (2024). Interactions of alpha-synuclein with membranes in Parkinson’s disease: Mechanisms and therapeutic strategies. Neurobiology of Disease, 201, 106646. 10.1016/j.nbd.2024.106646

Ma, M.-R., Hu, Z.-W., Zhao, Y.-F., Chen, Y.-X., & Li, Y.-M. (2016). Phosphorylation induces distinct alpha-synuclein strain formation. Scientific Reports, 6, 37130. 10.1038/srep37130

Machiya, Y., Hara, S., Arawaka, S., Fukushima, S., Sato, H., Sakamoto, M., Koyama, S., & Kato, T. (2010). Phosphorylated alpha-synuclein at Ser-129 is targeted to the proteasome pathway in a ubiquitin-independent manner. The Journal of Biological Chemistry, 285(52), Article 52. 10.1074/jbc.M110.141952

McFarland, N. R., Fan, Z., Xu, K., Schwarzschild, M. A., Feany, M. B., Hyman, B. T., & McLean, P. J. (2009). Alpha-synuclein S129 phosphorylation mutants do not alter nigrostriatal toxicity in a rat model of Parkinson disease. Journal of Neuropathology and Experimental Neurology, 68(5), 515–524. 10.1097/NEN.0b013e3181a24b53

Mizuguchi, H., Xu, Z., Ishii-Watabe, A., Uchida, E., & Hayakawa, T. (2000). IRES-dependent second gene expression is significantly lower than cap-dependent first gene expression in a bicistronic vector. Molecular Therapy: The Journal of the American Society of Gene Therapy, 1(4), 376–382. 10.1006/mthe.2000.0050

Muntané, G., Ferrer, I., & Martinez-Vicente, M. (2012). α-synuclein phosphorylation and truncation are normal events in the adult human brain. Neuroscience, 200, 106–119. 10.1016/j.neuroscience.2011.10.042

Neumann, M., Kahle, P. J., Giasson, B. I., Ozmen, L., Borroni, E., Spooren, W., M üller, V., Odoy, S., Fujiwara, H., & Hasegawa, M. (2002). Misfolded proteinase K – resistant hyperphosphorylated α-synuclein in aged transgenic mice with locomotor deterioration and in human α-synucleinopathies. The Journal of Clinical Investigation, 110 (10), 1429–1439.

Oueslati, A. (2016). Implication of Alpha-Synuclein Phosphorylation at S129 in Synucleinopathies: What Have We Learned in the Last Decade? Journal of Parkinson’s Disease, 6(1), Article 1. 10.3233/JPD-160779

Oueslati, A., Schneider, B. L., Aebischer, P., & Lashuel, H. A. (2013). Polo-like kinase 2 regulates selective autophagic α-synuclein clearance and suppresses its toxicity in vivo. Proceedings of the National Academy of Sciences of the United States of America, 110(41), Article 41. 10.1073/pnas.1309991110

Pelletier, J., & Sonenberg, N. (1988). Internal initiation of translation of eukaryotic mRNA directed by a sequence derived from poliovirus RNA. Nature, 334(6180), 320–325. 10.1038/334320a0

Polymeropoulos, M. H., Lavedan, C., Leroy, E., Ide, S. E., Dehejia, A., Dutra, A., Pike, B., Root, H., Rubenstein, J., & Boyer, R. (1997). Mutation in the α-synuclein gene identified in families with Parkinson ‘s disease. Science, 276 (5321), 2045–2047.

Ramalingam, N., Brontesi, L., Jin, S.-X., Selkoe, D. J., & Dettmer, U. (2023). Dynamic reversibility of α-synuclein serine-129 phosphorylation is impaired in synucleinopathy models. EMBO Reports, 24(12), Article 12. 10.15252/embr.202357145

Ramalingam, N., & Dettmer, U. (2023). α-Synuclein serine129 phosphorylation—The physiology of pathology. Molecular Neurodegeneration, 18(1), 84. 10.1186/s13024-023-00680-x

Ramalingam, N., Haass, C., & Dettmer, U. (2024). Physiological roles of α-synuclein serine-129 phosphorylation—Not an oxymoron. Trends in Neurosciences, 47(7), Article 7. 10.1016/j.tins.2024.05.005

Ramalingam, N., Jin, S.-X., Moors, T. E., Fonseca-Ornelas, L., Shimanaka, K., Lei, S., Cam, H. P., Watson, A. H., Brontesi, L., Ding, L., Hacibaloglu, D. Y., Jiang, H., Choi, S. J., Kanter, E., Liu, L., Bartels, T., Nuber, S., Sulzer, D., Mosharov, E. V., … Dettmer, U. (2023). Dynamic physiological α-synuclein S129 phosphorylation is driven by neuronal activity. NPJ Parkinson’s Disease, 9(1), Article 1. 10.1038/s41531-023-00444-w

San Gil, R., Berg, T., & Ecroyd, H. (2017). Using bicistronic constructs to evaluate the chaperone activities of heat shock proteins in cells. Scientific Reports, 7(1), 2387. 10.1038/s41598-017-02459-9

Spillantini, M. G., Schmidt, M. L., Lee, V. M., Trojanowski, J. Q., Jakes, R., & Goedert, M. (1997). Alpha-synuclein in Lewy bodies. Nature, 388(6645), Article 6645. 10.1038/42166

Su, D., Cui, Y., He, C., Yin, P., Bai, R., Zhu, J., Lam, J. S., Zhang, J., Yan, R., & Zheng, X. (2025). Projections for prevalence of Parkinson ‘s disease and its driving factors in 195 countries and territories to 2050: Modelling study of Global Burden of Disease Study 2021. Bmj, 388. Tenreiro, S.

Szymczak, A. L., Workman, C. J., Wang, Y., Vignali, K. M., Dilioglou, S., Vanin, E. F., & Vignali, D. A. A. (2004). Correction of multi-gene deficiency in vivo using a single ‘self-cleaving’ 2A peptide-based retroviral vector. Nature Biotechnology, 22(5), 589–594. 10.1038/nbt957

Tenreiro, S., Reimão-Pinto, M. M., Antas, P., Rino, J., Wawrzycka, D., Macedo, D., Rosado-Ramos, R., Amen, T., Waiss, M., Magalhães, F., Gomes, A., Santos, C. N., Kaganovich, D., & Outeiro, T. F. (2014). Phosphorylation modulates clearance of alpha-synuclein inclusions in a yeast model of Parkinson’s disease. PLoS Genetics, 10(5), Article 5. 10.1371/journal.pgen.1004302

Tripathi, A., Alnakhala, H., Brontesi, L., Selkoe, D., & Dettmer, U. (2024). RXR nuclear receptor signaling modulates lipid metabolism and triggers lysosomal clearance of alpha-synuclein in neuronal models of synucleinopathy. Cellular and Molecular Life Sciences: CMLS, 81(1), Article 1. 10.1007/s00018-024-05373-2

Wang, S., Xu, B., Liou, L.-C., Ren, Q., Huang, S., Luo, Y., Zhang, Z., & Witt, S. N. (2012). α-Synuclein disrupts stress signaling by inhibiting polo-like kinase Cdc5/Plk2. Proceedings of the National Academy of Sciences of the United States of America, 109(40), 16119–16124. 10.1073/pnas.1206286109

Yeboah, F., Kim, T.-E., Bill, A., & Dettmer, U. (2019). Dynamic behaviors of α-synuclein and tau in the cellular context: New mechanistic insights and therapeutic opportunities in neurodegeneration. Neurobiology of Disease, 132, 104543. 10.1016/j.nbd.2019.104543

Zarranz, J. J., Alegre, J., Gómez-Esteban, J. C., Lezcano, E., Ros, R., Ampuero, I., Vidal, L., Hoenicka, J., Rodriguez, O., Atarés, B., Llorens, V., Gomez Tortosa, E., del Ser, T., Muñoz, D. G., & de Yebenes, J. G. (2004). The new mutation, E46K, of alpha-synuclein causes Parkinson and Lewy body dementia. Annals of Neurology, 55(2), Article 2. 10.1002/ana.10795

