## Supplemental Figure 1 for "A Dual Assay to Compare Protein Levels and Toxicity of Alpha-Synuclein Variants: Acute Expression of Wild-Type versus S129A" (from another version of this preprint: version 1, 2025-09-23)

**Supplement**

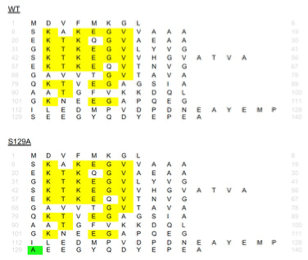

**Supplementary Figure 1:** Schematic of WT and S129A human αS with its amino acid sequence aligned via the KTKEGV motifs. Gray numbers on the sides denote the amino acid position. Amino acids that fully conform to ‘KTKEGV’ are highlighted in yellow. Alanine is highlighted in green.
